# Deep Learning Approach to Identifying Breast Cancer Subtypes Using High-Dimensional Genomic Data

**DOI:** 10.1101/629865

**Authors:** Runpu Chen, Le Yang, Steve Goodison, Yijun Sun

## Abstract

**Motivation:** Cancer subtype classification has the potential to significantly improve disease prognosis and develop individualized patient management. Existing methods are limited by their ability to handle extremely high-dimensional data and by the influence of misleading, irrelevant factors, resulting in ambiguous and overlapping subtypes.

**Results:** To address the above issues, we proposed a novel approach to disentangling and eliminating irrelevant factors by leveraging the power of deep learning. Specifically, we designed a deep learning framework, referred to as DeepType, that performs joint supervised classification, unsupervised clustering and dimensionality reduction to learn cancer-relevant data representation with cluster structure. We applied DeepType to the METABRIC breast cancer dataset and compared its performance to state-of-the-art methods. DeepType significantly outperformed the existing methods, identifying more robust subtypes while using fewer genes. The new approach provides a framework for the derivation of more accurate and robust molecular cancer subtypes by using increasingly complex, multi-source data.

**Availability and implementation:** An open-source software package for the proposed method is freely available at www.acsu.buffalo.edu/~yijunsun/lab/DeepType.html.

## 1 Introduction

Human cancer is a heterogeneous disease initiated by random somatic mutations and driven by multiple genomic alterations (Hanahan and Weinberg, 2011; Sun *et al*., 2017). In order to move towards personalized treatment regimes, cancers of specific tissues have been divided into subtypes based on the molecular profiles of primary tumors (Sørlie *et al*., 2001; Curtis *et al*., 2012; Parker *et al*., 2009). The premise is that patients of the same molecular subtypes are likely to have similar disease etiology, responses to therapy, and clinical outcomes. Thus, molecular subtyping can reveal information valuable for a range of cancer studies from etiology and tumor biology to prognosis and personalized medicine.

Most early work on molecular subtyping has been performed on data obtained from breast cancer tissues (Sørlie *et al*., 2001, 2003). Typically, breast cancer is not lethal immediately, and thus there is an opportunity to assist with prognostication and patient management using molecular information. Molecular subtyping of breast cancer initially focused on mRNA data obtained from microarray platforms and parsed molecular profiles to stratify patients according to clinical outcomes (Sørlie *et al*., 2001). Refinement of the subtype categories through validation in independent datasets identified five broad subtypes, including normal-like, luminal A, luminal B, basal, and HER2+, each with distinct clinical outcomes (Sørlie *et al*., 2003; Parker *et al*., 2009). These early studies completely altered our views of breast cancer and offered a foundation for the development of therapies tailored to specific subtypes. However, possibly due to the small number of tumor samples used in initial analyses and the technical limitations of the methods used for gene selection and clustering analysis, several large-scale benchmark studies have demonstrated that the current stratification of breast cancer is only approximate, and that the high degree of ambiguity in existing subtyping systems induces uncertainty in the classification of new patients (Weigelt *et al*., 2010; Mackay *et al*., 2011).

The desire for levels of accuracy that can ultimately lead to clinical utility continues to drive the field to refine breast cancer subtypes (Parker *et al*., 2009; Haibe-Kains *et al*., 2012; Shen *et al*., 2013; Sun *et al*., 2014, 2017) and to identify molecular subtypes in other cancers (Abeshouse *et al*., 2015). The recent establishment of international cancer genome consortia (Cancer Genome Atlas Network, 2012; Abeshouse *et al*., 2015; Curtis *et al*., 2012) has generally overcome the sample-size issue. In this paper, we focus mainly on developing methods to address the computational challenges associated with detecting cancer related genes and biologically meaningful subtypes using high-dimensional genomics data. Molecular subtyping can be formulated as a supervised-learning problem, that is, to use established tumor subtypes as class labels to perform gene selection and construct a model for the classification of new patients. However, as mentioned above, current subtyping systems provide only a rough stratification of cancer, and supervised-learning based approaches may not enable us to identify novel subtypes. This is because the primary goal of supervised learning is to identify genes to achieve the maximum separation of samples from different subtypes, and genes that support novel subtypes can be considered irrelevant and removed. Consequently, most existing methods were developed within the unsupervised-learning framework. Representative work includes SparseK (Witten and Tibshirani, 2010), iCluster (Shen *et al*., 2009, 2013) and non-negative matrix factorization (Kormaksson *et al*., 2012). A major issue with existing methods is that there is no guarantee that subtypes identified through *de novo* clustering are biologically relevant. Presumably, genomics data records all ongoing biological processes in a cell or tissue, where multiple factors interact with each other in a complex and entangled manner. Tumor samples can be grouped based on factors that are not related to the actual disease (e.g., race and eye color). A possible way to address the issue is to use previously established results to guide the detection of new subtypes. However, as the name suggests, *de novo* clustering completely ignores results from previous efforts. Another major limitation is that for computational considerations most existing methods perform data dimensionality reduction through linear transformation (e.g., feature weighting used in SparseK (Witten and Tibshirani, 2010)). Thus, they cannot adequately deal with complex non-linear data and extract pertinent information to detect subtypes residing in non-linear manifolds in a high-dimensional space. Finally, some existing methods do not scale well to handle high-dimensional data. For example, iCluster (Shen *et al*., 2009, 2013) involves matrix inversion and thus can only process a few thousands of genes. A commonly used practice is to perform preprocessing and retain only the most variant genes (Curtis *et al*., 2012). However, there is no guarantee that low-variant genes contain no information and the cut-offs used to select variant genes were usually set somehow arbitrarily.

The above observations motivated us to develop a novel deep-learning based approach, referred to as DeepType, that performs cancer subtyping through joint supervised and unsupervised learning but addresses their respective limitations. Due to the ability to learn good data representation, deep learning has recently achieved state-of-the-art performance in computer vision, pattern recognition and bioinformatics (LeCun *et al*., 2015; Zheng *et al*., 2019). For our purpose, by leveraging the power of a multi-layer neural network for representation learning, we map raw genomics data into a space where clusters can be easily detected. To ensure the biological relevance of detected clusters, we incorporate prior biological knowledge to guide representation learning. We train the neural network by minimizing a unified objective function consisting of a classification loss, a clustering loss and a sparsity penalty. The training process can be easily performed by using a mini-batch gradient descent method. Thus, our method can handle large datasets with extremely high dimensionality. Although the idea of using deep learning for clustering is not new (see, e.g., Xie *et al.* (2016)), to the best of our knowledge, this work represents the *first* attempt to use deep learning to perform joint supervised and unsupervised learning for cancer subtype classification. A large-scale experiment was performed that demonstrated that DeepType significantly outperformed the existing approaches. The new approach provides a framework for the derivation of more accurate and robust molecular cancer subtypes by using increasingly complex genomic data.

## 2 Methods

In this section, we present a detailed description of the proposed method for cancer subtype identification. We also propose novel procedures for optimizing the associated objective function and estimating the hyper-parameters.

### 2.1 Deep Learning for Cancer Subtype Identification

Let **X** = [**x**_1_, …, **x**_*N*_] be a cohort of tumor samples and **Y** = [**y**_1_, …, **y**_*N*_] be a rough stratification of the samples (e.g., subtyping results from previous studies), where **x**_*n*_ ∈ ℝ^*D*^ is the *n*-th sample and **y**_*n*_ ∈ ℝ^*J*^ is the corresponding class label vector with *y*_*jn*_ = 1 if **x**_*n*_ belongs to the *j*-th group and 0 otherwise. Our goal is to identify a small set of cancer related genes and perform clustering analysis on the detected genes to refine existing classification systems and detect novel subtypes. To this end, we utilize the representation power of a multi-layer neural network to project raw data onto a representation space where clusters can be easily detected. As discussed above, clusters identified through unsupervised learning may not be biologically relevant. To address the issue, we impose an additional constraint that the detected clusters are concordance with previous results. Specifically, we cast it as a supervised-learning problem, that is, to find a representation space where the class labels can be accurately predicted.

Figure 1 depicts the network structure of the proposed method. It consists of an input layer, *M* hidden layers, a classification layer and a clustering module. The *M*-th hidden layer is designated as the representation layer, the output of which is fed into the classification layer and the clustering module. Mathematically, the neural network can be described as follows:

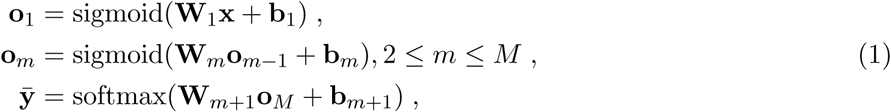

where **W**_*m*_, **b**_*m*_, and **o**_*m*_ are the weight matrix, bias term and output of the *m*-th layer, respectively, and 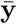 is the output of the classification layer. For the purpose of this study, we use sigmoid and softmax as the activation functions for the hidden and classification layers, respectively. For notational convenience, let 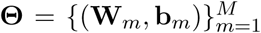 and denote 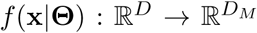 as the mapping function that projects raw data onto a representation space, where *D*_*M*_ is the number of the nodes in the representation layer and *D*_*M*_ ≪ *D*.

**Figure 1:**
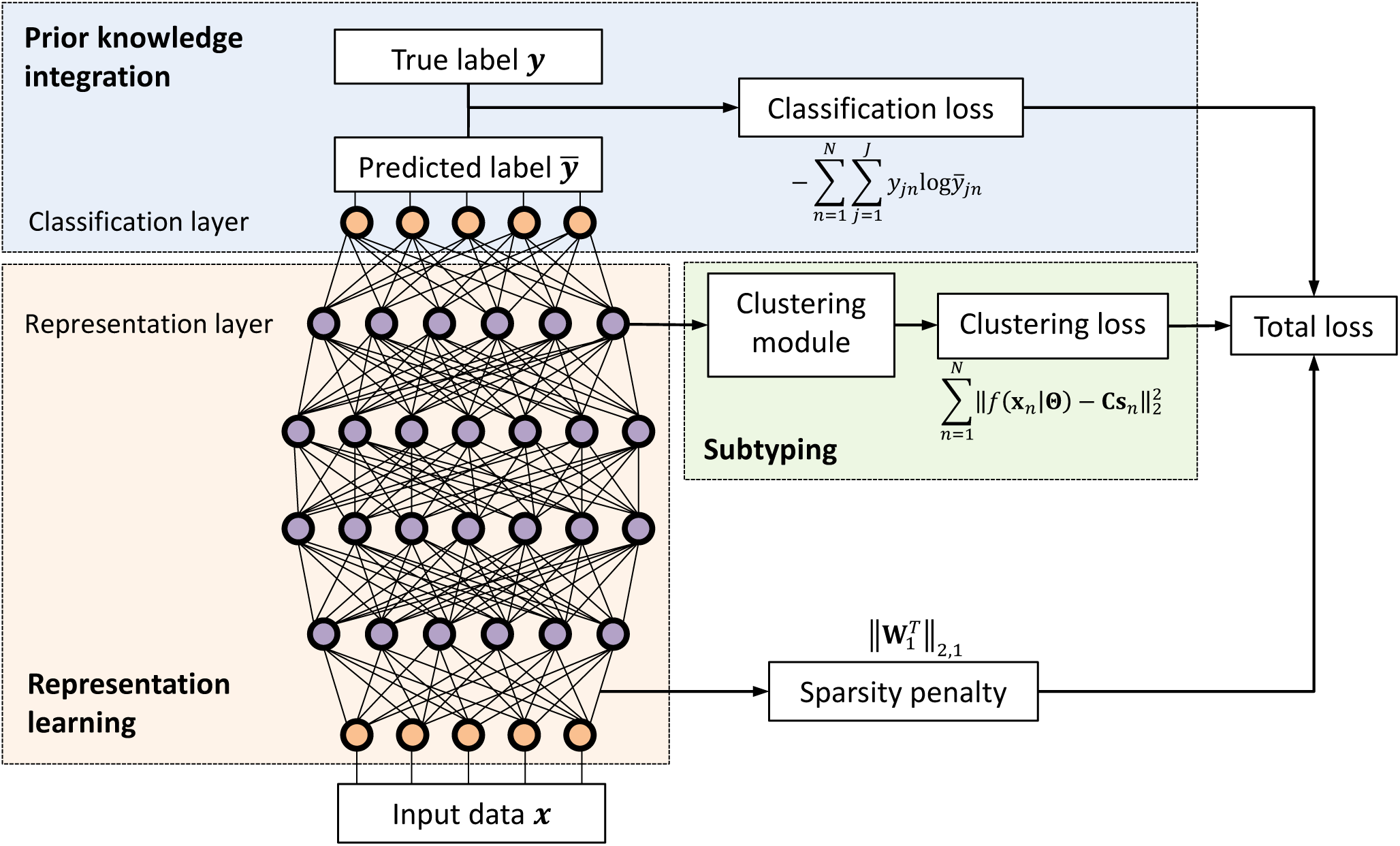
Overview of the proposed deep learning based method for cancer molecular subtyping. It consists of three major components: representation learning, prior knowledge integration, and subtyping. The first part maps raw genomics data onto a representation space, the second part incorporates prior biological knowledge to guide representation learning, and the third part generates subtyping results. The network parameters are learned by minimizing a unified objective function consisting of a classification loss, a clustering loss and a sparsity penalty.

We optimize network parameters **Θ** through joint supervised and unsupervised learning by minimizing an objective function that consists of a classification loss, a clustering loss and a regularization term. The classification loss measures the discrepancy between the predicted and given class labels. By construction, the *j*-th element of 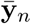 can be interpreted as the probability of **x**_*n*_ belonging to the *j*-th group. Thus, we use the cross entropy to quantify the classification loss:

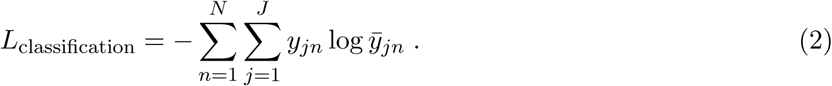

We use the *K*-means method (Lloyd, 1982) to detect clusters in the representation space. The loss function optimized by *K*-means is given by

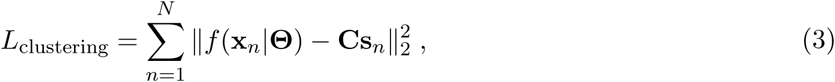

subject to 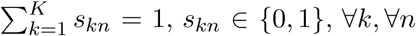, where *K* is the number of clusters, **C** is a center matrix with each column representing a cluster center, and **s**_*n*_ is a binary vector where *s*_*kn*_ = 1 if **x**_*n*_ is assigned to cluster *k* and 0 otherwise. Finally, we impose an *ℓ*_2,1_-norm regularization (Nie *et al*., 2010) on the weight matrix of the first layer to control the model complexity and to select cancer related genes:

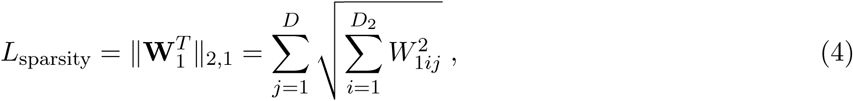

where *W*_1*ij*_ is the *ij*-th element of **W**_1_ and *D*_2_ is the number of the nodes in the second layer. The *ℓ*_2,1_-norm regularization has an effect of automatically determining the number of nodes activated in the input layer, and thus the number of genes used in downstream subtyping analysis.

Combining the above three losses, we obtain the following novel formulation for cancer subtype identification:

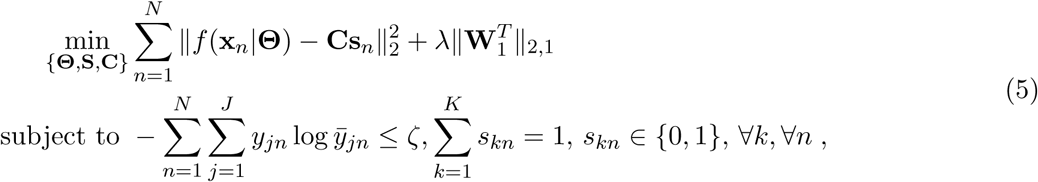

where **S** = [**s**_1_, …, **s**_*N*_] and *λ* is a regularization parameter that controls the sparseness of weight matrix **W**_1_. The above formulation can be interpreted as finding a representation space to minimize the clustering loss while maintaining the classification loss smaller than a user defined upper bound *ζ*. For ease of optimization, we move the classification-loss constraint to the objective function and write the problem in the following equivalent form:

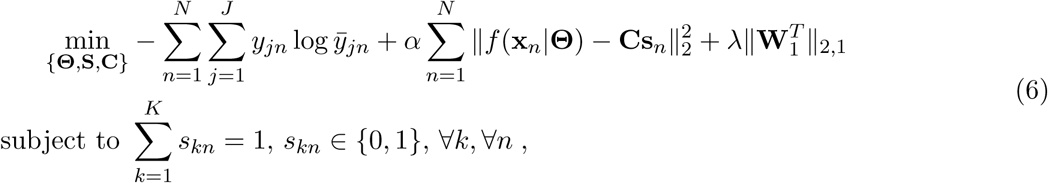

where *α* is a tradeoff parameter that controls the balance between the classification and clustering performance. In the following sections, we describe how to solve the above optimization problem and estimate the hyper-parameters.

### 2.2 Optimization

The above optimization problem contains three sets of variables, namely, network parameters **Θ**, assignment matrix **S**, and cluster centers **C**. It is difficult to solve the problem directly since the parameters are coupled and **S** is a binary matrix. To address the issue, we partition the variables into two groups, i.e., **Θ** and (**S, C**), and employ an alternating optimization strategy to solve the problem. Specifically, we first perform pre-training to initialize the network by ignoring the clustering module (i.e., setting *α* = 0).

Then, we fix **Θ** and transform the problem into

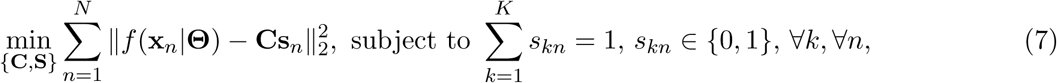

which can be readily solved by using the standard *K*-means method. Then, we fix (**S, C**) and write the problem as

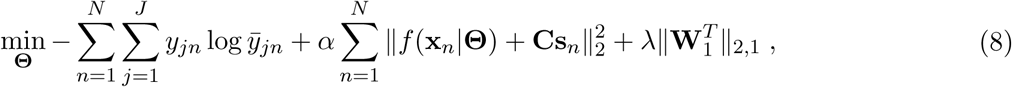

which can be optimized through back-propagation by using the mini-batch based stochastic gradient descent method (Kingma and Ba, 2014). The above procedures iterate until convergence.

### 2.3 Parameter Estimation

We describe how to estimate the three hyper-parameters of the proposed method, namely regularization parameter *λ*, tradeoff parameter *α*, and number of clusters *K*. In order to avoid a computationally expensive three-dimensional grid search, we first ignore the clustering module by setting *α* = 0 and perform supervised learning to estimate *λ*. The rationale is that previous subtyping results could provide us with sufficient information to determine the value of *λ*. Specifically, we randomly partition training data into ten equally-sized sub-datasets, perform ten-fold cross-validation and estimate *λ* by using the one-standard-error rule (Hastie *et al*., 2009). Once we determine the value of *λ*, we perform *K*-means analysis on the outputs of the representation layer and pre-estimate the number of clusters, denoted as 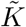, as the one that maximizes the average silhouette width (Wiwie *et al*., 2015). Since the data representation is obtained through supervised learning, which tends to group samples with the same labels together, 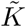 is likely to be the lower bound of the true value. Let 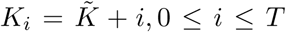. For each *K*_*i*_, we train a deep-learning model by using different *α* values and record the corresponding ten-fold cross-validation classification errors. By design, *α* controls the tradeoff between the classification and clustering performance, and the classification error increases with the increase of *α*. Again, by using the one-standard-error rule, for each *K*_*i*_, we find the largest *α*, denoted as *α*_*i*_, that results in a classification error that is within one standard deviation of the one obtained by setting *α* = 0 (i.e., we require that the obtained classifier does not perform significantly worse than the existing subtyping system), and record the corresponding average silhouette width *s*_*i*_. Once we run over all possible *K*_*i*_, we obtain *T* + 1 triplets 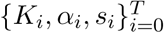. Finally, we determine the number of clusters *K* and the tradeoff parameter *α* as the pair that yields the largest average silhouette width. The pseudo-code of the proposed procedure is given in Algorithm S1, and the proposed procedure performed quite well in our numerical experiment (see Figure S1).

## 3 Experiments

We conducted a large-scale experiment on breast and bladder cancers to demonstrate the effectiveness of the proposed method. Due to space limit, here we report only the results of the breast cancer study and present the bladder cancer results in Supplementary Data.

### 3.1 Experiment Setting

The breast cancer dataset was obtained from the METABRIC study (Curtis *et al*., 2012), which contains the expression profiles of 25,160 genes from 1,989 primary breast tumor samples and 144 normal breast tissue samples. It is probably the largest single breast cancer dataset assayed to date. For computational convenience, we retained only the top 20,000 most variant genes for the downstream analysis. For model construction and performance evaluation, we randomly partitioned the data into a training and test datasets, containing 80% and 20% of the samples, respectively. In this study, we used the PAM50 subtypes (Parker *et al*., 2009) as class labels in the training process. We designed a four-layer neural network for the joint supervised and unsupervised learning. The numbers of the nodes in the input layer, the two hidden layers and the output layer were set to 20,000, 1,024, 512, and 6, respectively. We employed the Adam method (Kingma and Ba, 2014) to tune the parameters of the model. The learning rate was set to 1e-3, the numbers of training epochs for model initialization and the joint supervised and unsupervised training were set to 300 and 1,500, respectively, and the batch size was set to 256. By using the method proposed in Section 2.3, the number of clusters *K*, the tradeoff parameter *α* and the regularization parameter *λ* were estimated to be 11, 1.2 and 0.006, respectively (see Figure S1). To ensure that the constructed model did not overfit the data, we tracked the training and validation losses in the training process (see Figure S2), and no sign of over-fitting was observed.

### 3.2 Clinically Relevant Subtypes Revealed by DeepType

By applying the proposed method to the breast cancer dataset, a total of 218 genes were selected and 11 clusters were detected. To visualize the identified clusters, we applied t-SNE (van der Maaten and Hinton, 2008) to the outputs of the representation layer. Figure 2(a-b) present the sample distributions of the identified clusters and their PAM50 compositions, respectively. We can see that nearly all of the normal tissue samples were grouped into a single cluster (i.e., Cluster 0), and the tumor samples were grouped into ten well-separated clusters, labelled as DeepType 1-10. To demonstrate the clinical relevance of the identified tumor subtypes, a disease-specific survival data analysis was performed. Figure 2(c) shows that the ten subtypes were associated with distinct prognostic outcomes (logrank test, *p*-value *<* 1.22e-19). Further internal and external validation analysis of the detected clusters is presented in Sections 3.3 and 3.4.

**Figure 2:**
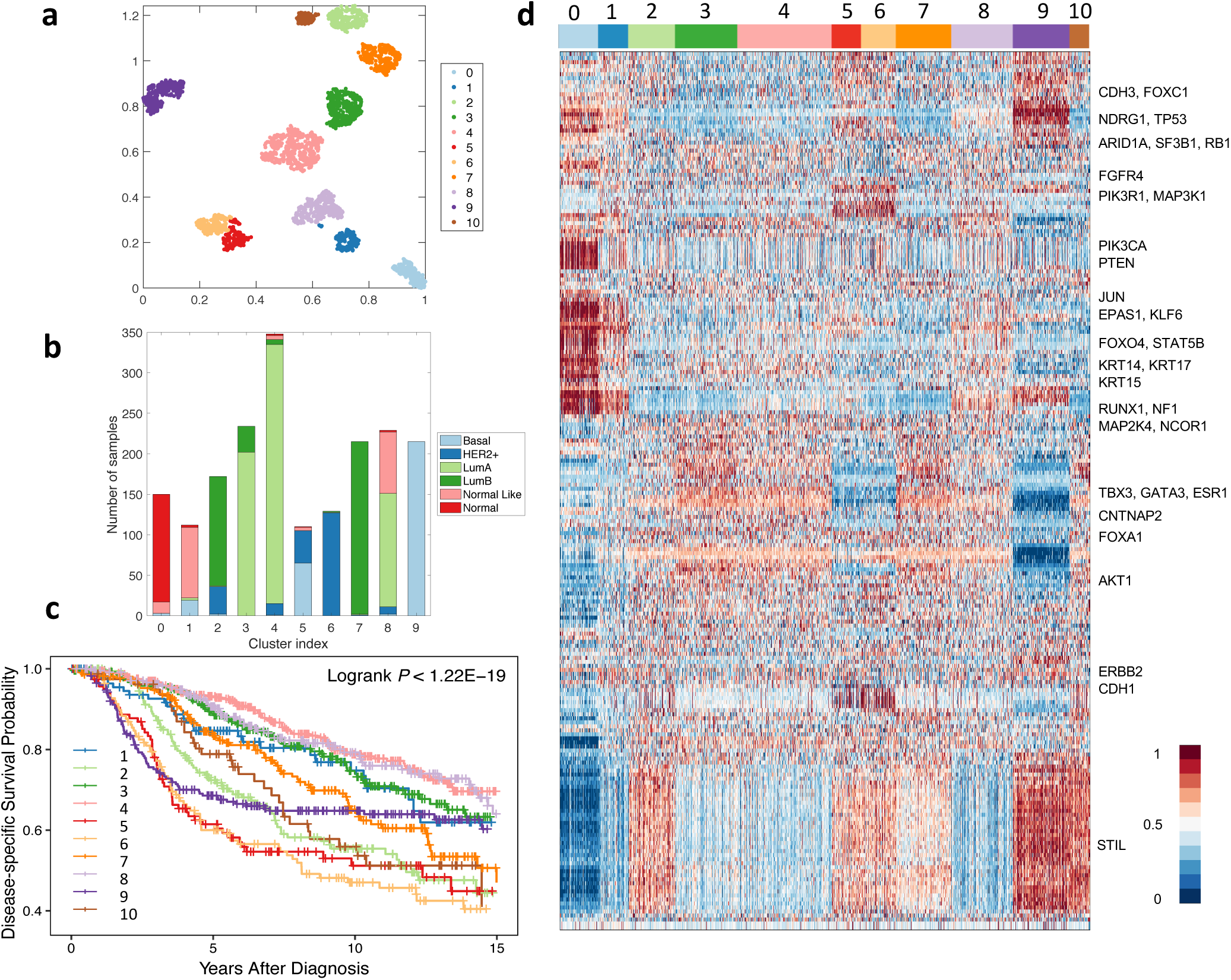
DeepType identified ten clinically relevant breast cancer subtypes. (a) The sample distributions of the identified clusters visualized by t-SNE. Nearly all of the normal tissue samples were grouped into a single cluster (i.e., Cluster 0), and the tumor samples were grouped into ten well-separated clusters, labelled as DeepType 1-10. (b) The PAM50 composition of the identified clusters. (c) Survival data analysis showed that the ten identified subtypes were associated with distinct clinical outcomes. (d) The heatmap of the 218 selected genes showed that the identified clusters exhibited distinct transcriptional characteristics on several gene modules. The samples were arranged by the clustering assignments, and the expression levels were linearly scaled into [0, 1] across samples.

Figure 2(d) represents the heatmap of the 218 selected genes. The descriptions of the genes are given in Table S1. The detected subtypes contain distinct transcriptional characteristics associated with several gene co-expression modules and key cancer genes. Most normal-like samples were grouped into DeepType 1, and have an expression pattern similar to normal samples. The luminal A samples were divided into DeepTypes 3, 4 and 8 with low expression on the *STIL* module (key gene: *STIL*) and intermediate expression on the *GATA3* module (key genes: *TBX3, GATA3, ESR1, CNTNAP2* and *FOXA1*). Among the three subtypes, the expression of the *KRT* family (key genes: *KRT14, KRT15* and *KRT17*) were highest in DeepType 8, intermediate in DeepType 4, and lowest in DeepType 3. The luminal B samples were partitioned into DeepTypes 2, 7 and 10, with intermediate to high expression of the *GATA3* and *STIL* gene modules, and low expression of *CDH3* and *FOXC1*. Among the three subtypes, the expression of the genes in the *STIL* module was highest in DeepType 10, intermediate in DeepType 2 and lowest in DeepType 7. DeepTypes 5 and 6, which were dominated by mixed HER2+/basal and HER2+ samples, respectively, had very high expression on *ERBB2* and *CDH1* and low expression on *TBX3, GATA3* and *ESR1* genes. DeepType 9, composed entirely of basal samples, had low expression in the *GATA3* module and high expression in the *STIL* and *KRT* modules. The distinct expression patterns and prognostic outcomes of the detected clusters suggest that the proposed method is able to detect new breast cancer subtypes beyond the PAM50 classification, and a further analysis could reveal information of the breast cancer molecular taxonomy in a higher level of resolution.

### 3.3 Comparison Study

To further demonstrate the effectiveness of the proposed method, we compared it with two state-of-the-art methods, namely SparseK (Witten and Tibshirani, 2010) and iCluster (Shen *et al*., 2009). Both methods perform feature selection and clustering analysis simultaneously, and iCluster was also used in the METABRIC study (Curtis *et al*., 2012). The source code of the two methods was downloaded from the CRAN website^1^,^2^. Following Shen *et al.* (2013), *we tuned the parameters of iCluster (i*.*e*., *the number of clusters K* and the sparsity penalty coefficient *λ*) by maximizing the reproducibility index. SparseK also contains two parameters, the number of clusters *K* and the *ℓ*_1_ regularization parameter *λ*. By using the method described in Witten and Tibshirani (2010), we first estimated the optimal *λ* for each *K*, and then determined the value of the optimal *K* based on gap statistic (Tibshirani *et al*., 2001). To test the ability of the three methods to handle high-dimensional data, we generated four datasets each containing a different number of the most variant genes, ranging from 5,000, 10,000, 15,000 and 20,000. Although we herein considered only gene expression data, it is possible to perform cancer subtyping by integrating genomics data from different platforms. Therefore, the ability to handle high-dimensional data is an important consideration in algorithm development. Below, we performed a series of quantitative and qualitative analyses to compare the performance of the three methods.

We first visualized the sample distributions of the clusters detected by the three methods (Figure 3). Since iCluster failed on the datasets with 15,000 and 20,000 genes due to the need of performing matrix inversion of high-dimensional data, we considered only the results generated by using the dataset with 10,000 genes. We can see that DeepType identified eleven well-defined clusters, nearly all normal tissue samples were grouped into a single cluster, and the clusters that composed of tumor samples were well-separated and highly concordant with the PAM50 labels. In contrast, for SparseK and iCluster, the normal tissue samples were grouped into multiple clusters, which suggests that genes unrelated to cancer were selected. Moreover, the tumor samples with different PAM50 labels overlapped considerably, and did not exhibit a clear clustering structure.

**Figure 3:**
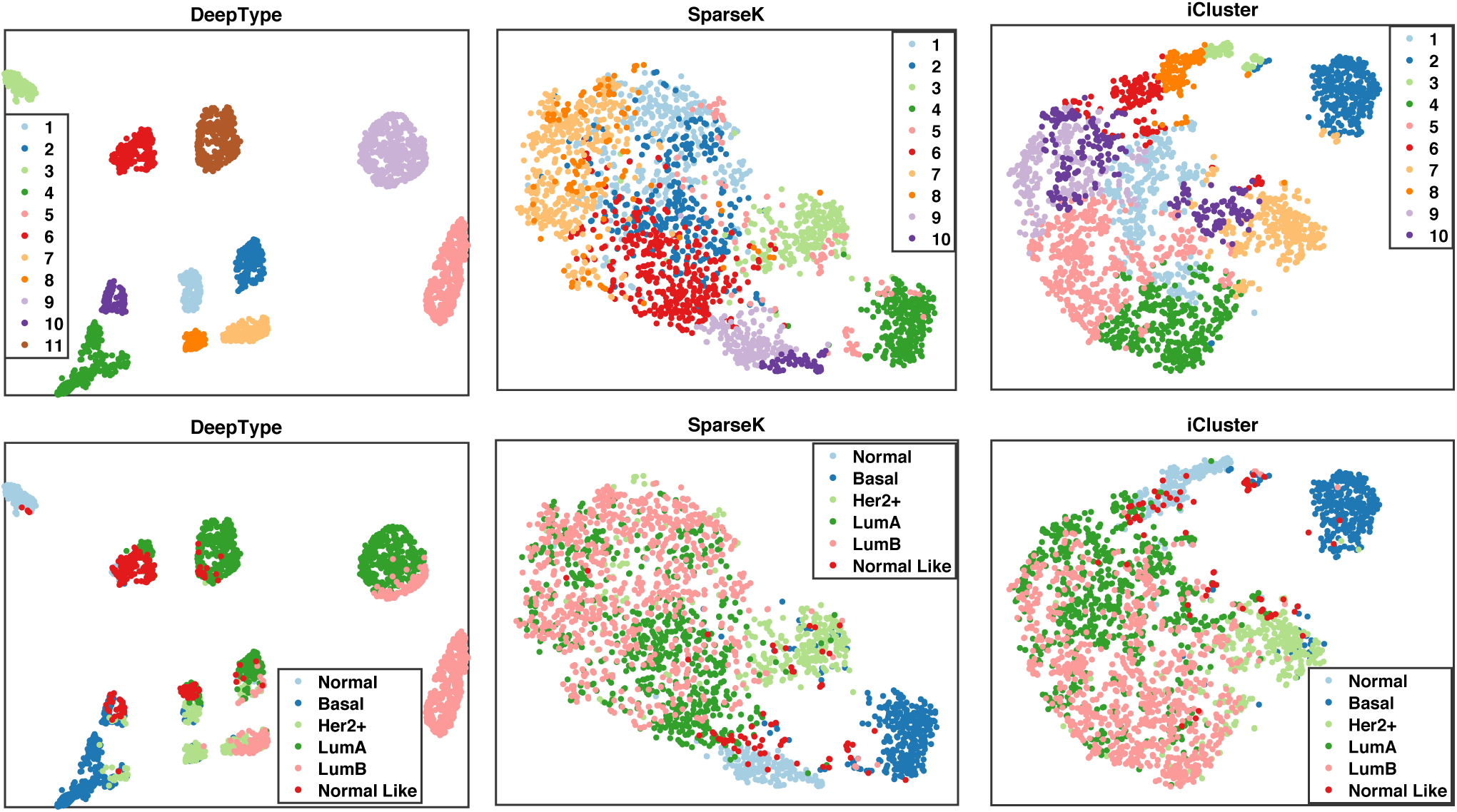
Visualization of the sample distributions of the clusters detected by three methods applied to data containing 10,000 most variant genes. Each sample was color-coded by its clustering assignment (top) and PAM50 label (bottom). DeepType revealed a clear eleven-cluster structure including a cluster comprising primarily normal tissue samples.

We then performed a series of external and internal evaluations of the clusters detected by the three methods. For external evaluation, we assessed the concordance between the identified cancer subtypes and some widely used clinical and prognostic characteristics of breast cancer, including the PAM50 subtype (Parker *et al*., 2009), histological grade, Nottingham prognostic index (NPI) (Haybittle *et al*., 1982), gene expression grade index (GGI) (Sotiriou *et al*., 2006) and the Oncotype DX prognostic test (Sparano *et al*., 2018) (see Table S2 for a detailed description). Specifically, we used average purity and normalized mutual information (NMI) to evaluate the extent to which the identified subtypes matched the above described characteristics. The results are reported in Table 1. Our analysis showed that the subtypes identified by DeepType were highly concordant with the clinical variables and prognostic information. In all cases, the results generated by DeepType matched the PAM50 labels to the highest degree. This is expected since the PAM50 labels were used in training DeepType. Our method also produced the highest agreement with the histological grades, NPI and GGI. Notably, when compared with Oncotype DX, the average purities and NMI scores of DeepType were much higher than the other two methods. This is highly significant since while both NPI and GGI provide some values in predicting the clinical outcomes of breast cancer patients, Oncotype DX is the only test supported by level II evidence (Sparano *et al*., 2018). We performed a Wilcoxon rank-sum test to compare the overall performance of DeepType and the two competing methods. The *p*-values are 7.7e-14 (DeepType vs. SparseK) and 1.3e-19 (DeepType vs. iCluster).

**Table 1:** External evaluation of subtypes identified by three methods applied to datasets with a various number of input genes. iCluster failed on datasets with 15,000 and 20,000 genes. DeepType significantly outperformed SparseK (*p*-value ≤ 7.7e-14) and iCluster (*p*-value ≤ 1.3e-19, Wilcoxon rank-sum test).

|  |  | Average Purity |  |  |  | NMI |  |  |  |
| --- | --- | --- | --- | --- | --- | --- | --- | --- | --- |
|  |  | 5000 | 10000 | 15000 | 20000 | 5000 | 10000 | 15000 | 20000 |
| PAM50 | DeepType | **0.86** | **0.80** | **0.85** | **0.87** | **0.62** | **0.56** | **0.62** | **0.61** |
|  | SparseK | 0.65 | 0.68 | 0.64 | 0.63 | 0.39 | 0.40 | 0.37 | 0.38 |
|  | iCluster | 0.43 | 0.65 | / | / | 0.08 | 0.34 | / | / |
| Histological Grade | DeepType | **0.67** | **0.67** | **0.66** | **0.67** | **0.12** | **0.12** | **0.12** | **0.13** |
|  | SparseK | 0.63 | 0.66 | 0.65 | 0.63 | 0.11 | 0.10 | **0.12** | 0.09 |
|  | iCluster | 0.55 | 0.63 | / | / | 0.04 | 0.11 | / | / |
| NPI | DeepType | **0.57** | 0.55 | **0.60** | **0.58** | **0.08** | **0.09** | **0.10** | **0.07** |
|  | SparseK | 0.56 | 0.55 | 0.59 | 0.58 | 0.07 | **0.09** | 0.07 | **0.07** |
|  | iCluster | 0.56 | **0.57** | / | / | 0.04 | **0.09** | / | / |
| GGI | DeepType | **0.69** | 0.68 | **0.70** | **0.69** | **0.15** | **0.16** | **0.14** | **0.13** |
|  | SparseK | **0.69** | **0.70** | 0.70 | **0.69** | 0.12 | 0.14 | 0.13 | 0.12 |
|  | iCluster | 0.68 | 0.67 | / | / | 0.04 | 0.11 | / | / |
| Oncotype DX | DeepType | **0.88** | **0.85** | **0.87** | **0.86** | **0.25** | **0.26** | **0.27** | **0.24** |
|  | SparseK | 0.75 | 0.78 | 0.78 | 0.76 | 0.14 | 0.16 | 0.15 | 0.13 |
|  | iCluster | 0.64 | 0.74 | / | / | 0.06 | 0.13 | / | / |

We next performed internal evaluation of the subtypes identified by the three methods. Internal evaluation utilizes only the intrinsic information of cluster assignments to assess the quality of obtained clusters, and compactness and separability are the two most important considerations (Halkidi *et al*., 2001). A compact and separable clustering structure means that samples in each cluster are homogeneous and different clusters are far away from each other, allowing new patients to be assigned with high certainty and low ambiguity. For the purpose of this study, we used the silhouette width (Wiwie *et al*., 2015) and the Davies–Bouldin index (Davies and Bouldin, 1979) to quantify the cluster compactness and separability. The results are reported in Table 2. In all cases, DeepType resulted in the highest silhouette width and the lowest Davies–Bouldin index, which is consistent with the visualization result presented in Figure 3. To compare the overall performance, the Wilcoxon rank-sum test was performed. Deeptype significantly outperformed SparseK (*p*-value ≤ 7.8e-5) and iCluster (*p*-value ≤ 7.8e-5). Our analysis suggested that our method resulted in subtypes with significantly higher cluster quality than the competing methods.

**Table 2:** Internal evaluation of subtypes identified by three methods applied to datasets with a various number of input genes. The Davies–Bouldin index is a value in [0, inf), and a smaller value suggests a better clustering scheme. DeepType significantly outperformed SparseK (*p*-value ≤ 7.8e-5) and iCluster (*p*-value ≤ 7.8e-5, Wilcoxon rank-sum test).

|  | Silhouette width |  |  | Davies–Bouldin index |  |  |
| --- | --- | --- | --- | --- | --- | --- |
|  | DeepType | SparseK | iCluster | DeepType | SparseK | iCluster |
| 5000 | **0.48** | 0.17 | 0.33 | **1.01** | 1.88 | 1.79 |
| 10000 | **0.48** | 0.22 | 0.33 | **0.87** | 1.94 | 1.23 |
| 15000 | **0.44** | 0.19 | / | **0.69** | 1.92 | / |
| 20000 | **0.63** | 0.15 | / | **0.67** | 2.31 | / |

Finally, we compared the ability of the three methods to select relevant genes from high-dimensional data for clustering analysis. Table 3 reports the numbers of genes selected by the three methods applied to the data with a various number of input genes. Notably, while DeepType achieved the best result in terms of both internal and external criteria, it selected the fewest genes in all cases. For clinical applications, the ability to select fewer genes can help to develop a more economic clinical assay for breast cancer subtype identification.

**Table 3:** The numbers of genes selected by DeepType, iCluster and SparseK applied to datasets containing a various number of input genes.

| # of input genes | DeepType | SparseK | iCluster |
| --- | --- | --- | --- |
| 5000 | **182** | 949 | 521 |
| 10000 | **239** | 982 | 728 |
| 15000 | **250** | 918 | / |
| 20000 | **218** | 886 | / |

### 3.4 Validation Study

To demonstrate the generalization capability of the proposed method, we performed a validation study using the METABRIC data for training and SUPERTAM data (Haibe-Kains *et al*., 2012) for testing. The SUPERTAM dataset contains the expression profiles of 13,092 genes from 856 breast tumor samples. Prior to the analysis, we identified 10,087 genes present in both datasets and used ComBat (Johnson *et al*., 2007) to remove batch effects. Using the expression measures of the selected genes, we trained a deep-learning model using the METABRIC dataset and identified eleven clusters including one comprising dominantly normal samples. We then applied the constructed model to the validation dataset and classified each sample into one of the eleven clusters using the nearest shrunken centroid classifier (Tibshirani *et al*., 2002). Since the SUPERTAM data does not contain normal samples, only three samples were classified into the normal cluster and thus omitted in the further analysis. Figure 4(a-d) presents the sample distributions and PAM50 compositions of the identified clusters. We observed that the clusters detected in the two datasets were compact and well-separated and had similar PAM50 compositions. To provide a quantitative analysis of the reproducibility of the detected clusters, we employed the strategy proposed in Kapp and Tibshirani (2006) and calculated the in-group proportion (IGP) score and *p*-value for each cluster (Figure 4(e)). Our analysis showed that the identified clusters were reproducible (*p*-value = 0) and that the proposed method generalizes well on independent datasets.

**Figure 4:**
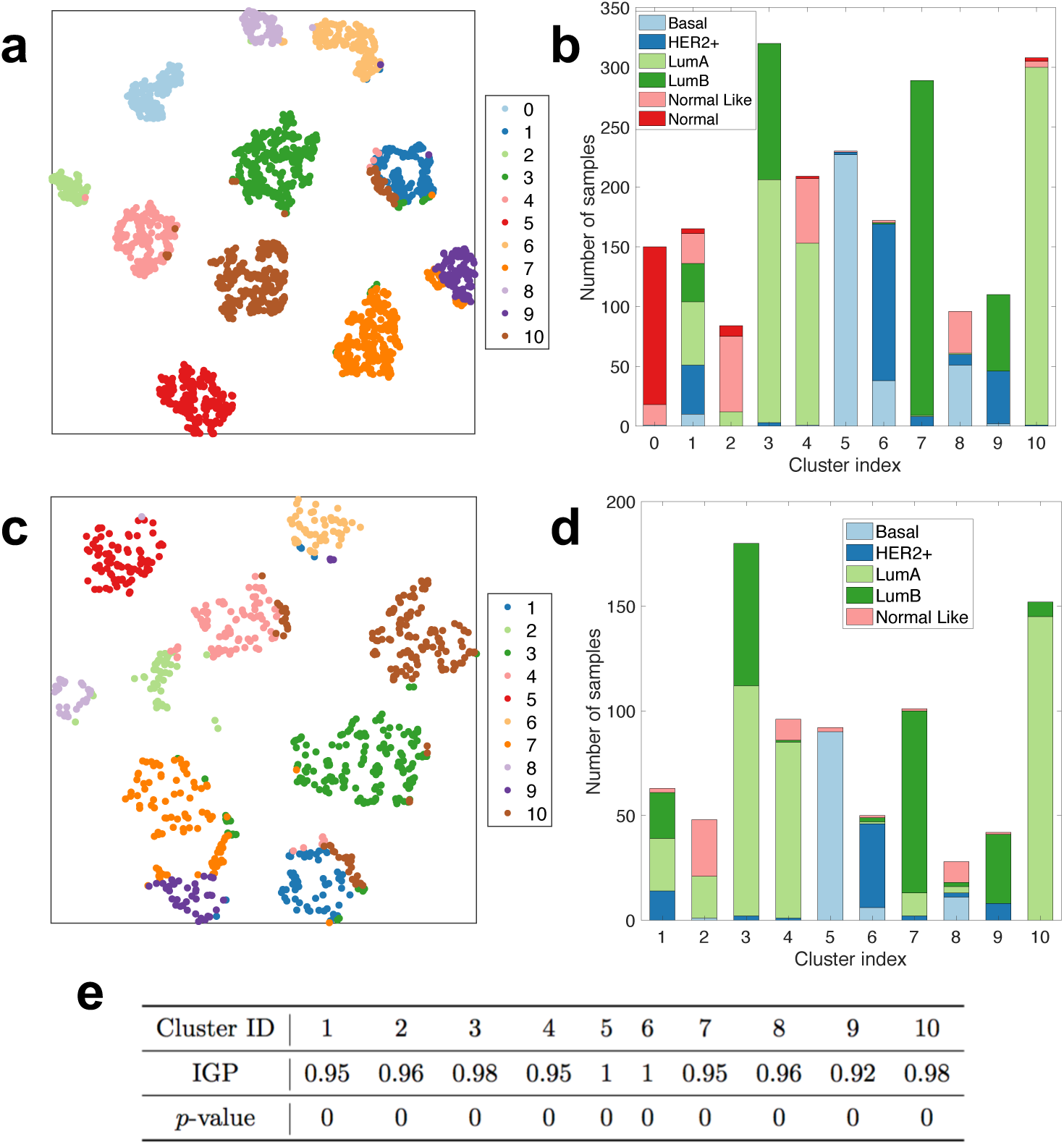
Results of a validation study performed on the METABRIC and SUPERTAM datasets. (a-d) The clusters detected in the METABRIC (top row) and SUPERTAM datasets (middle row) were compact and well-separated and had very similar PAM50 compositions. (e) In-group proportion (IGP) scores and *p*-values (computed based on 1,000 permutations) showed that the clusters identified in the METABRIC data were reproducible in the SUPERTAM data.

### 3.5 Robustness Analysis

DeepType detects disease molecular subtypes through joint supervised and unsupervised learning, where the class labels from previous studies are usually error-prone. To investigate how DeepType performs in the presence of label noise, we performed a robustness analysis where we corrupted the PAM50 labels of a certain percentage of randomly selected samples in the METABRIC training dataset, constructed a deep-learning model using the corrupted data, and applied the model to the test dataset. To assess the performance of the constructed model, we computed the Rand index by comparing the cluster assignments of the test samples with their PAM50 labels and those obtained by using the original training dataset (i.e., no corrupted labels). To remove random variations, the experiment was repeated five times. Figure S3 presents the results obtained by using the training data containing a varying percentage of corrupted labels ranging from 0% to 20%. We can see that DeepType performed similarly with up to 10% label errors. Considering that the PAM50 label set itself contains an unknown percentage of errors, our method is very robust against label noise.

## 4 Discussion

In this paper, we developed a deep-learning based approach for cancer subtype identification that addresses some technical limitations of existing methods. The new method performed significantly better than two commonly used approaches in terms of both internal and external evaluation criteria. By leveraging the power of deep learning, the new method is able to handle data with extremely high dimensionality. We further demonstrated that the method generalizes well on independent datasets and is very robust against label noise.

The proposed method has several limitations that are worthwhile to mention. Usually, training a deep-learning model requires a large amount of data. The method is thus not applicable to cancers for which only a small number of samples have been assayed. In this study, we applied the method to breast and bladder cancers where molecular subtypes are well established and thus can be used to guide the detection of new subtypes. However, for many other cancers, molecular subtypes have not yet been well established. It is possible to use other clinical variables (e.g., tumor grade) to guide the identification of cancer subtypes and we have showed that our approach performed well in the presence of label noise. Further investigations are warranted to explore such possibilities.

In this paper, we presented a proof-of-concept study considering only gene expression data. Several studies have recently demonstrated that combining cross-platform data could provide more information for cancer subtype identification (see, e.g., Shen *et al.* (2013) and Zhang *et al.* (2012)). It is possible to use deep learning to integrate genomics data from different platforms, including mRNA, copy number, somatic mutation and methylation, for cancer subtyping. However, currently there are debates on how to design a network to process multiple data types (Wang *et al*., 2015). As the future work, we will perform a large-scale experiment to look into this issue to identify the optimal network structure for genomics data analysis. It is expected that more accurate and robust cancer subtypes would be revealed.

## Supplementary Data

### Algorithm 1 Hyper-parameter estimation (X; Y; 𝒜; ℒ; *T*)

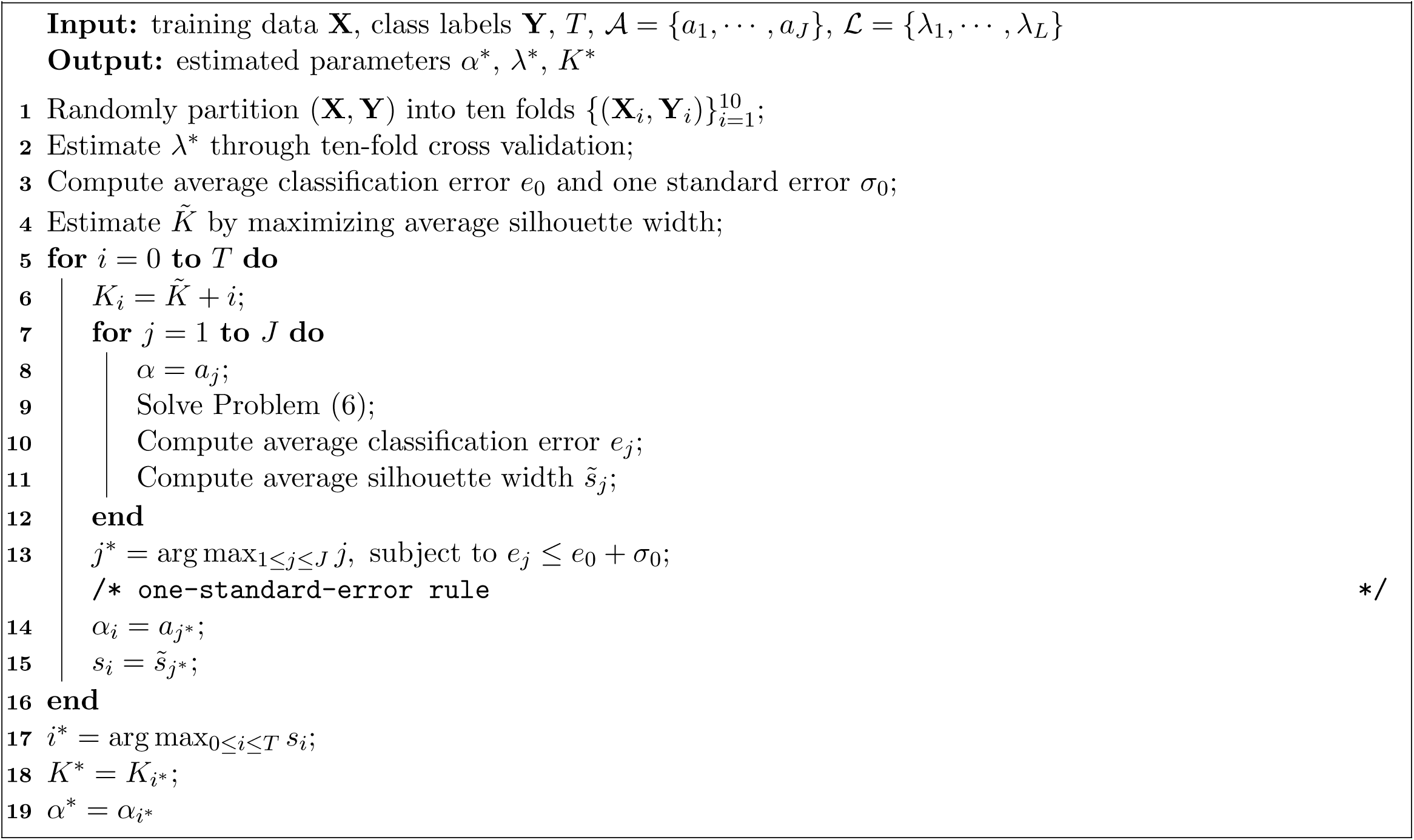

**Table S1:**
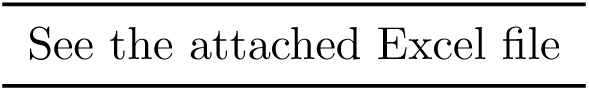
DeepType identified 218 genes to be informative for breast cancer subtyping See the attached Excel file

**Table S2:** Clinical and prognostic characteristics of breast cancer

| Characteristics | Class label |
| --- | --- |
| PAM50 subtype | basal, HER2+, luminal A/B, normal-like |
| Histological grade | 1, 2, 3 |
| NPI | 1, 2, 3, 4 |
| GGI | low risk, high risk |
| Oncotype DX | low risk, intermediate risk, high risk |

**Table S3:**
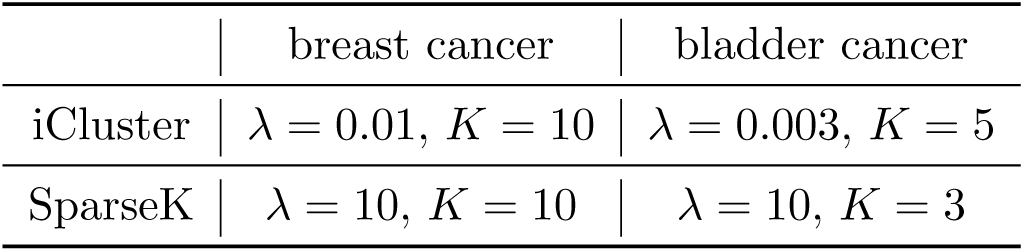
Parameters of iCluster and SparseK used in the breast and bladder cancer experiments

|  | breast cancer | bladder cancer |
| --- | --- | --- |
| iCluster | $\lambda = 0.01, K = 10$ | $\lambda = 0.003, K = 5$ |
| SparseK | $\lambda = 10, K = 10$ | $\lambda = 10, K = 3$ |

**Table S4:**
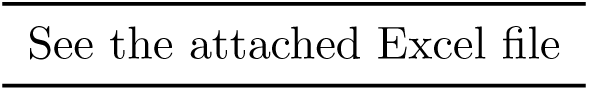
The indexes of the samples in the training and test datasets used in the breast cancer experiment (for the reproducibility purpose)

**Figure S1:**
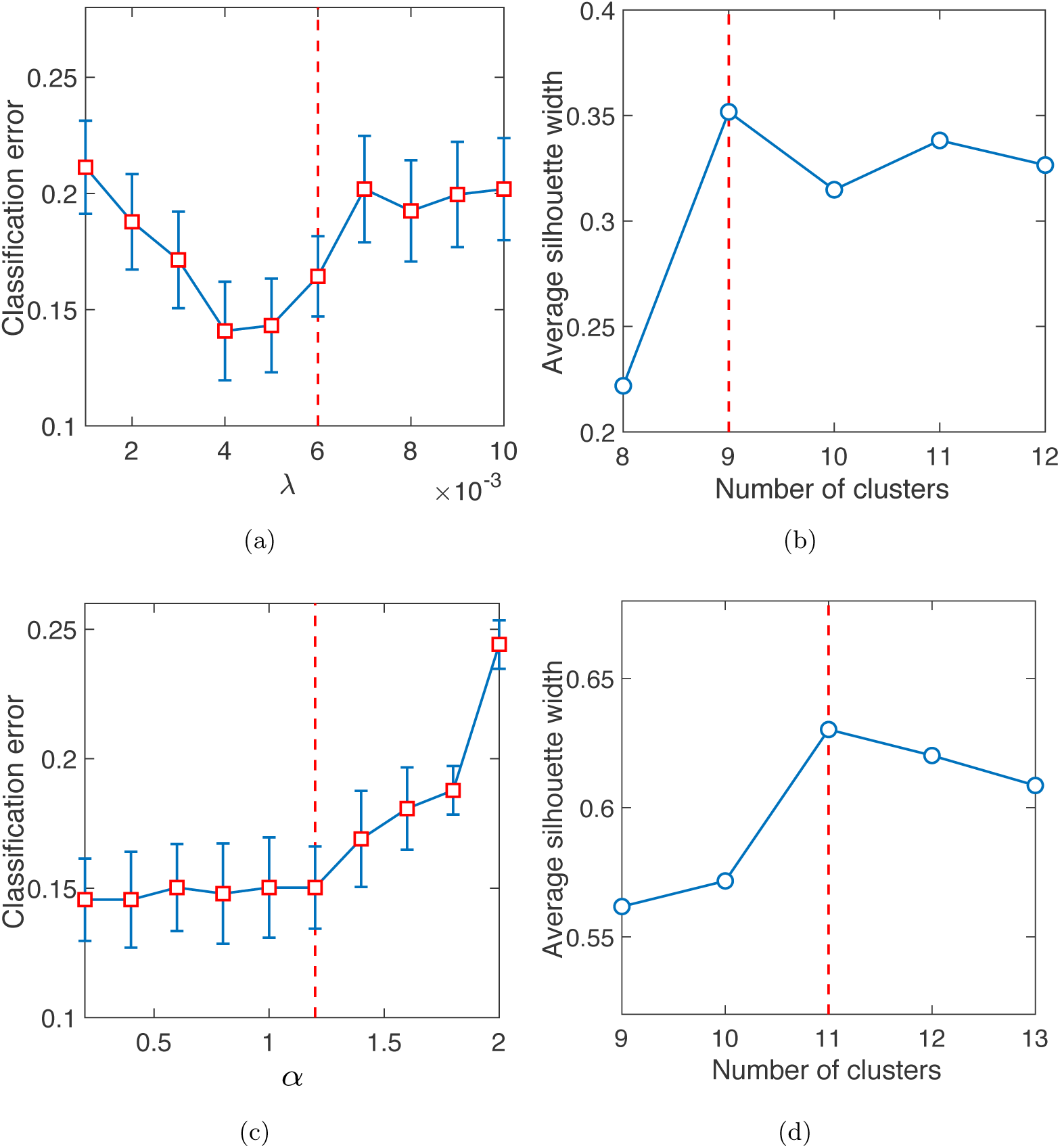
Hyper-parameter estimation. (a) The regularization parameter *λ* was estimated to be 0.006 based on the one-standard-error rule. (b) The number of clusters was pre-estimated to be 9 based on the average silhouette width. (c) We searched a range of values to estimate the number of clusters. For each *K* ≥ 9, we trained a deep learning model by using different *α* values and estimated the optimal *α* by using the one-standard-error rule. The figure presents an example showing that the optimal *α* was estimated to be 1.2 for *K* = 11. (d) The number of clusters was finally determined to be 11 by maximizing the average silhouette width. See Section 2.3 for a detailed description of hyper-parameter estimation.

**Figure S2:**
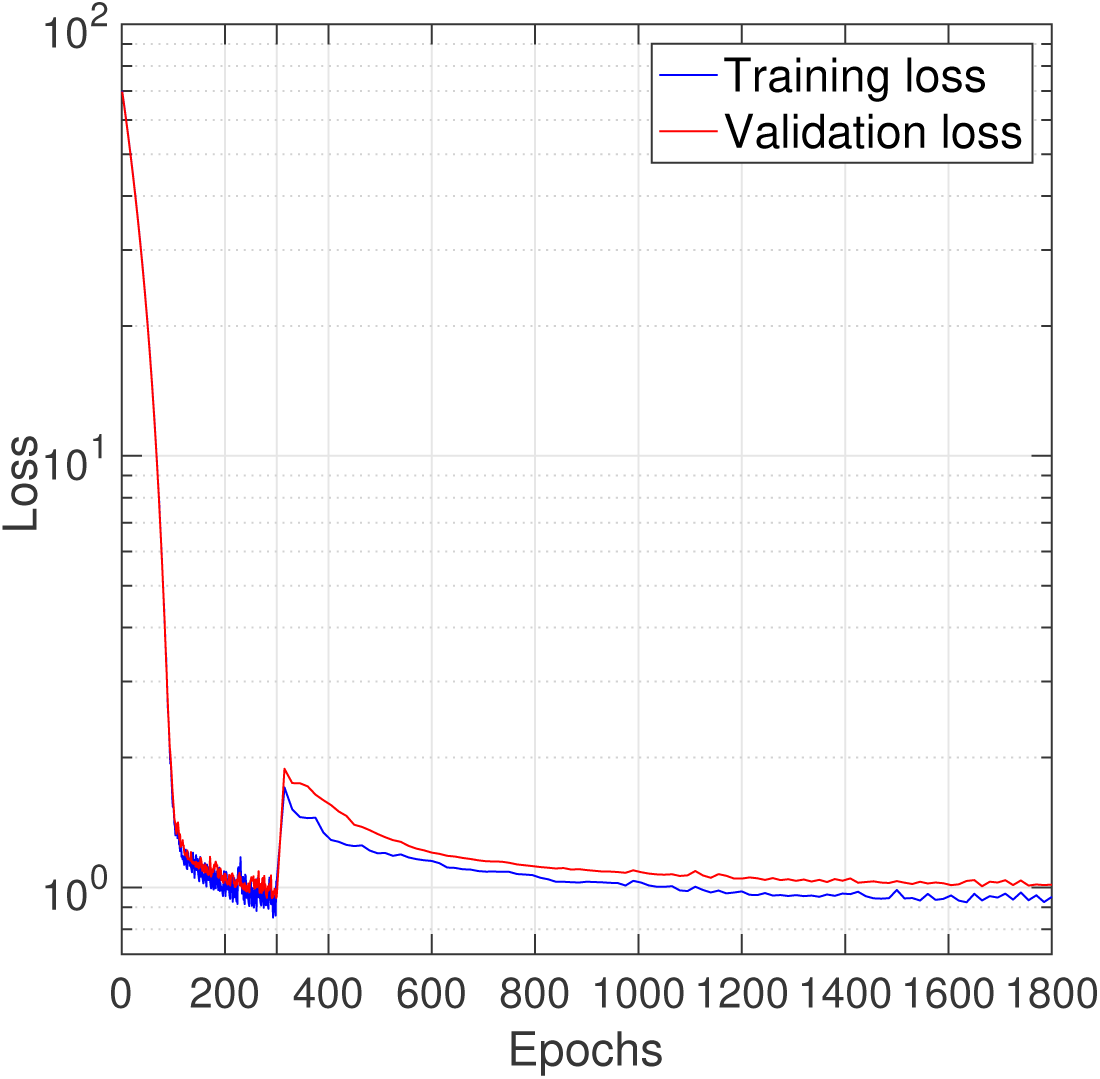
Curves of training and validation losses vs. epochs. No sign of over-fitting was observed. The first 300 epochs are for model initialization based on supervised learning and the remaining 1500 epochs are for model optimization based on joint supervised and unsupervised learning.

**Figure S3:**
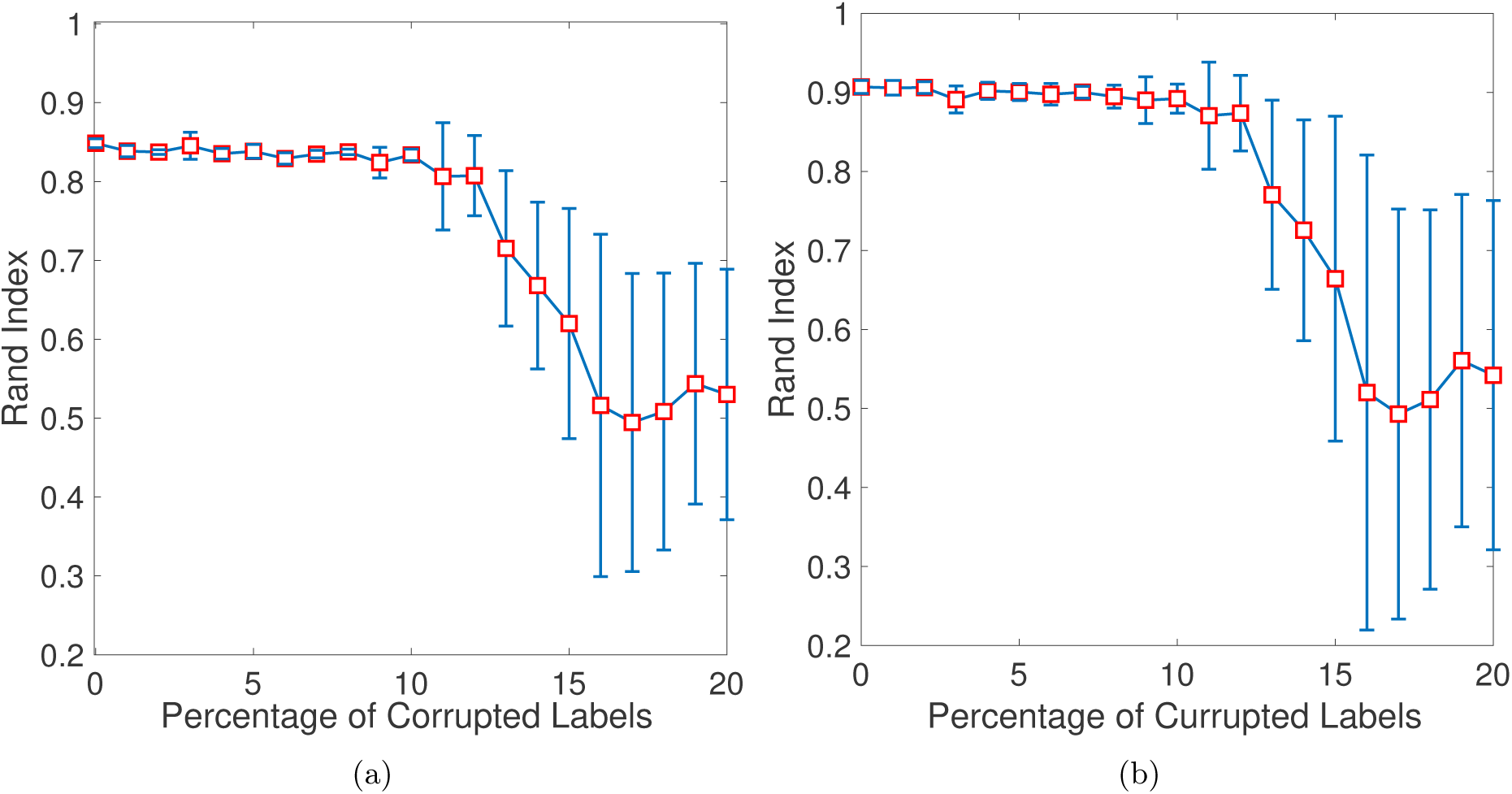
Rand indexes computed by comparing the cluster assignments of the test samples with their PAM50 labels (a) and those obtained by using the original training dataset (b). DeepType performed similarly when up to 10% of the class labels of the training data were corrupted.

### S1 Bladder Cancer Study

#### S1.1 Experiment Setting

The bladder cancer dataset was obtained from the TCGA project, which contains the expression profiles of 20,241 genes from 427 bladder tumor samples. We classified each tumor sample into one of the two UNC subtypes (Damrauer *et al*., 2014), namely luminal and basal, using the R package BLCAsubtyping (Kamoun *et al*., 2019), and used the UNC subtypes as the class labels. For model construction and performance evaluation, we randomly partitioned the data into a training and test dataset, containing 80% and 20% of the samples, respectively. Since the sample size is small, in order to avoid possible overfitting, we used only the top 10,000 most variant genes, and designed a three-layer neural network with 10,000, 32 and 2 nodes in the input layer, the hidden layer and the output layer, respectively. The number of clusters *K*, the tradeoff parameter *α* and the regularization parameter *λ* were estimated to be 4, 0.005 and 0.002, respectively. Other experiment settings were similar to those used in the breast cancer study.

#### S1.2 Clinically Relevant Subtypes Revealed by DeepType

By applying DeepType to the bladder cancer dataset, a total of 156 genes were selected and 4 clusters were detected. The descriptions of the selected genes are given in Table S5. To visualize the detected clusters, we applied t-SNE (van der Maaten and Hinton, 2008) to the outputs of the representation layer. Figure S4(a-b) presents the sample distributions of the identified clusters and their UNC-subtype compositions, respectively. We can see that the tumor samples were grouped into four well-defined clusters, including two luminal dominated clusters (labeled as luminal 1 and 2) and two basal dominated clusters (labeled as basal 1 and 2). To demonstrate the clinical relevance of the identified tumor subtypes, a survival data analysis was performed. Figure S4(c) shows that the four subtypes are associated with distinct prognostic outcomes (logrank test, *p*-value *<* 0.0001). Further internal and external validation analysis is presented in Section S1.3.

Figure S4(d) presents the heatmap of the 156 selected genes. We can clearly see two modules, one containing key genes *MSN, TNC* and *MUC16*, and the other containing key genes *TOX3* and *PDX1*. The difference in the gene expressions in the two modules divided the samples into two broad categories, i.e., basal and luminal. Specifically, the basal samples have high expressions in the *MSN* module and low expressions in the *TOX3* module, and the luminal samples are on the contrary. Within each UNC subtype, luminal 1 has higher expressions in the *MSN* module than luminal 2, while basal 1 has higher expressions in the *TOX3* module than basal 2. Using *t*-test and the Benjamini-Hochberg procedure (Benjamini and Hochberg, 1995), we identified 99 genes differentially expressed between luminal 1 and luminal 2, and 112 genes between basal 1 and basal 2 (FDR ≤ 0.05). Our analysis showed that DeepType is able to identify novel bladder cancer subtypes beyond the UNC subtyping system that are associated with distinct expression patterns.

#### S1.3 Comparison Study

For comparison, we applied SparseK and iCluster to the bladder cancer dataset. The parameters of the two methods were estimated in the same way as that used in the breast cancer study. We first visualized the sample distributions of the clusters detected by the three methods (Figure S5). As with the breast cancer study, we can see that the clusters identified by DeepType are much more compact than those detected by SparseK and iCluster. Then, we performed a series of external and internal evaluations of the quality of the identified clusters. For external evaluation, we assessed the concordance between cluster assignments and the UNC subtypes, the tumor pathological stages, and the risk of tumor recurrence computed based on a three-gene signature proposed in (Liu *et al*., 2017) (see Table S6 for a detailed description). For internal evaluation, we used the silhouette width and Davies-Bouldin index to quantify the compactness and separateness of the obtained clusters. The results are reported in Tables S7 and S8. In terms of external criteria, our method performed significantly better than SparseK and slightly better than iCluster. In terms of internal criteria, DeepType resulted in the highest silhouette width and the lowest Davies–Bouldin index, which is consistent with the visualization result presented in Figure S5. Our analysis suggested that our method resulted in subtypes with significantly higher cluster quality than the competing methods.

**Figure S4:**
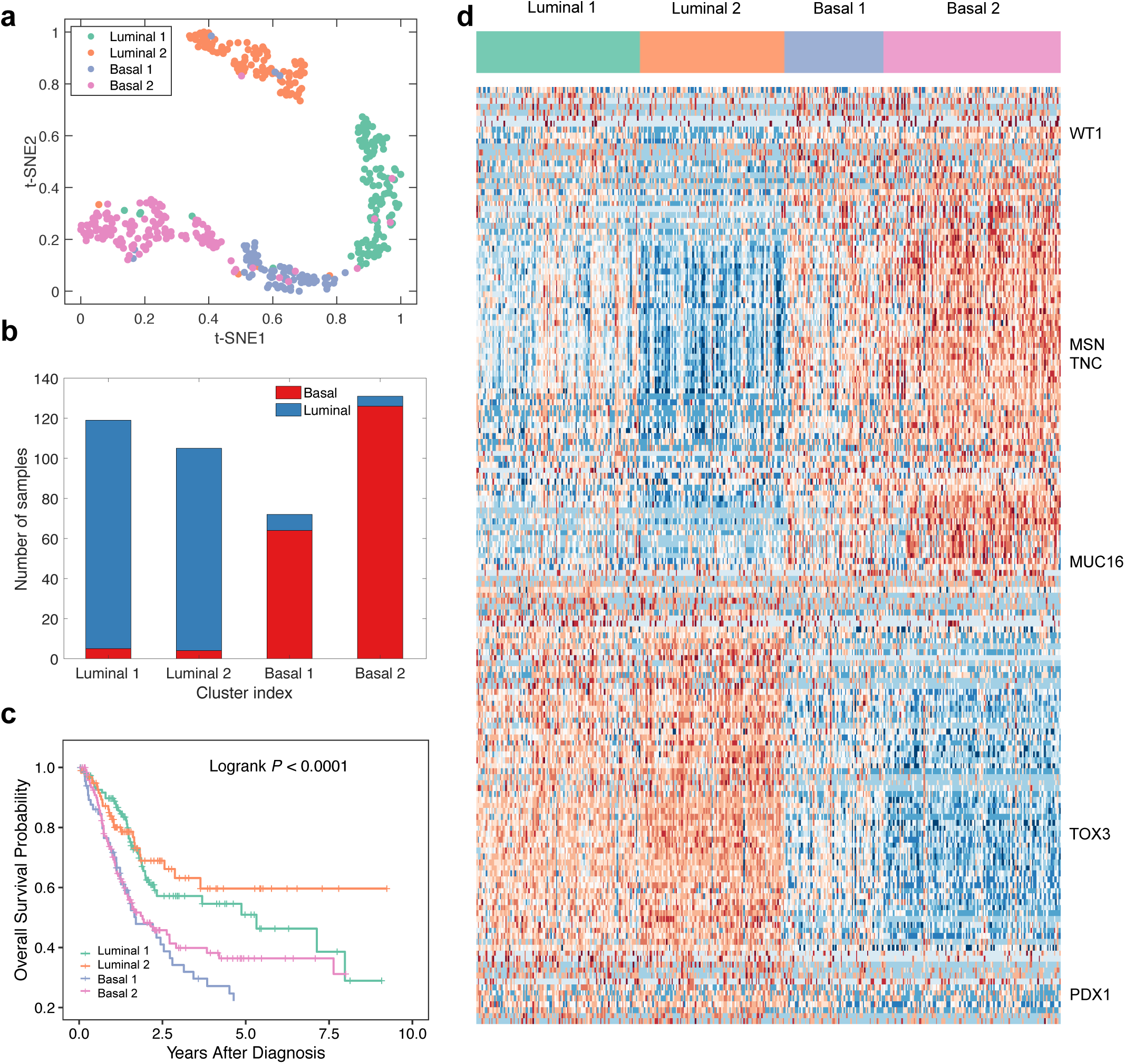
DeepType identified four clinically relevant bladder cancer subtypes. (a) The sample distribution of the identified clusters visualized by t-SNE. (b) The UNC-subtype composition of the four clusters. (c) The four identified subtypes are associated with distinct clinical outcomes. (d) The heatmap of the 156 selected genes showed that the identified clusters exhibited distinct transcriptional characteristics on several gene modules and key genes. The samples were arranged by the clustering assignments, and the expression data are linearly normalized into the scale of [0, 1] across samples.

**Figure S5:**
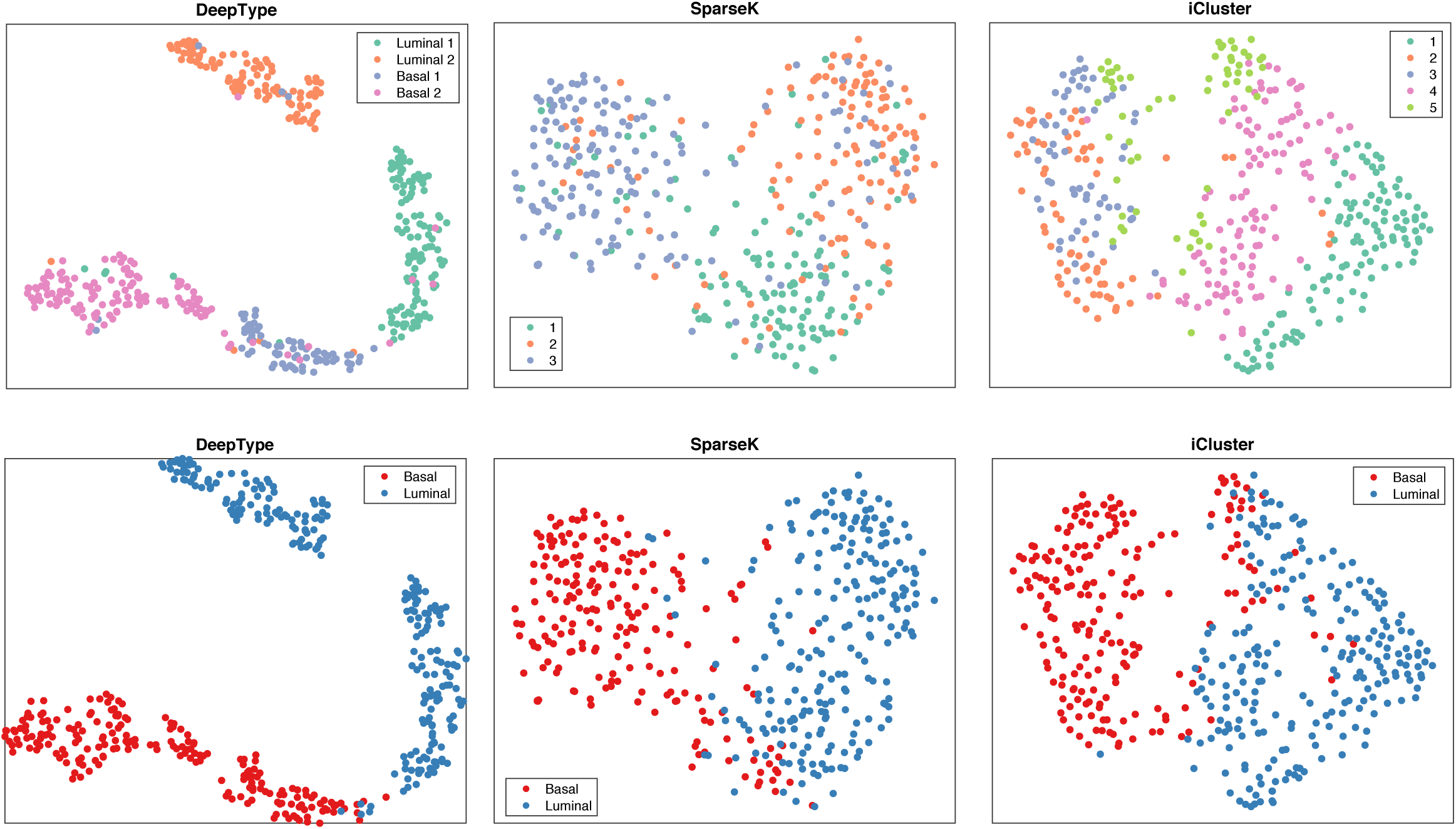
Visualization of the sample distributions of the clusters detected by three methods applied to the bladder cancer dataset. Each sample is color-coded by its clustering assignment (top) and UNC-subtype label (bottom). DeepType revealed a clear four-cluster structure.

**Table S5:**
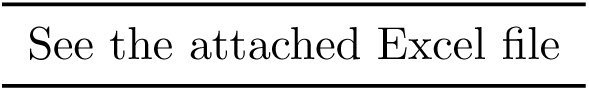
DeepType identified 156 genes to be informative for bladder cancer subtyping See the attached Excel file

**Table S6:**
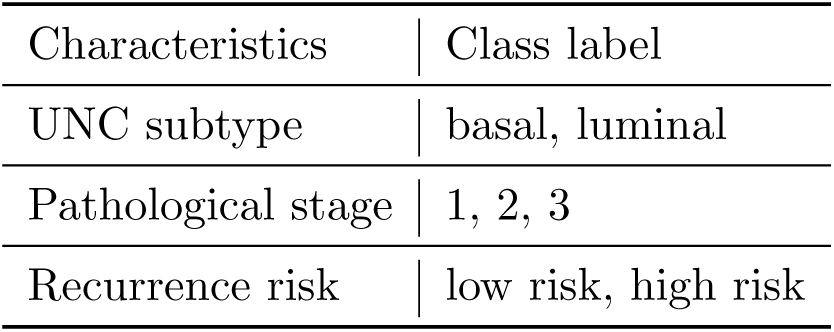
Clinical and prognostic characteristics of bladder cancer

| Characteristics | Class label |
| --- | --- |
| UNC subtype | basal, luminal |
| Pathological stage | 1, 2, 3 |
| Recurrence risk | low risk, high risk |

**Table S7:** External evaluation of subtypes identified by three methods applied to the bladder cancer dataset. DeepType significantly outperformed SparseK (*p* ≤ 0.001) and iCluster (*p* ≤ 0.03, Wilcoxon rank-sum test).

|  | Average Purity |  |  | NMI |  |  |
| --- | --- | --- | --- | --- | --- | --- |
|  | DeepType | SparseK | iCluster | DeepType | SparseK | iCluster |
| UNC subtypes | **0.95** | 0.75 | 0.91 | **0.51** | 0.16 | 0.43 |
| Pathological stage | **0.45** | 0.42 | 0.44 | **0.04** | 0.02 | **0.04** |
| Recurrence risk | **0.78** | 0.66 | 0.75 | **0.20** | 0.08 | **0.20** |

**Table S8:** Internal evaluation of subtypes identified by three methods applied to the bladder cancer dataset. The Davies-Bouldin index is a value in [0, inf), and a smaller value suggests a better clustering scheme. DeepType outperforms the competing methods by a large margin.

|  | DeepType | SparseK | iCluster |
| --- | --- | --- | --- |
| Silhouette width | **0.45** | 0.19 | 0.04 |
| Davias-Bouldin index | **1.00** | 2.56 | 3.29 |

## Footnotes

1 https://cran.r-project.org/web/packages/iCluster/index.html

2 https://cran.r-project.org/web/packages/sparcl/index.html

